# Allele-specific multi-sample copy number segmentation

**DOI:** 10.1101/166017

**Authors:** Edith M. Ross, Kerstin Haase, Peter Van Loo, Florian Markowetz

## Abstract

**Motivation:** Allele-specific copy number alterations are commonly used to trace the evolution of tumours. A key step of the analysis is to segment genomic data into regions of constant copy number. For precise phylogenetic inference, breakpoints shared between samples need to be aligned to each other.

**Results:** Here we present `asmultipcf`, an algorithm for allele-specific segmentation of multiple samples that infers private and shared segment boundaries of phylogenetically related samples. The output of this algorithm can directly be used for allele-specific copy number calling using ASCAT.

**Availability:** `asmultipcf` is available as part of the ASCAT R package (version 2.5) from github.com/Crick-CancerGenomics/ascat

## 1 Introduction

Allele-specific copy number alterations (CNAs) are commonly used to trace the evolution of a tumour. One of the most frequently used algorithms to infer these copy number changes is AS-CAT (Van Loo et al., 2010). It segments each sample separately and due to the noise in the data the inferred locations of shared breakpoints are likely to differ between samples. These differences can impair the analysis of phylogenetic relationships between the samples as it depends on the assumption that shared breakpoints appear at exactly the same location. To address this problem, Schwarz et al. (2015) performed extensive experimental break point validation, an expensive approach which has often been omitted by similar papers. Mangiola et al. (2016), for example, used a size-based heuristic filter for CNAs instead.

To rigorously address the problem of multi-sample breakpoint detection, we have developed `asmultipcf` (allele-specific multi-sample piecewise constant fitting), an algorithm performing allele-specific segmentation of multiple samples for the inference of private and shared segment boundaries of phylogenetically related samples. asmultipcf enforces joint segment boundaries across samples unless the data contains significant evidence that the breakpoints differ.

## 2 Approach

`asmultipcf` is based on the copy number segmentation algorithms developed by Nilsen et al. (2012), which use penalized least square principles to fit piecewise constant segments to the data and which are used by ASCAT (Van Loo et al., 2010) to infer allele specific copy numbers. asmpultipcf combines two algorithms by Nilsen et al. (2012), aspcf (an allele-specific single-sample segmentation method) and multipcf (a non-allele-specific multi-sample segmentation method), to enable the joint segmentation of multiple related samples in an allele specific manner. Additionally, `asmpultipcf` handles missing values, making extensive data filtering unnecessary.

### Input data

For each sample the following input data are required across germline heterozygous sites: (i) log ratios (logR), representing log-transformed copy numbers derived from sequencing depth or SNP array data, and (ii) B allele frequencies (BAF), describing the allelic imbalance of SNPs. Both measurements can be derived from whole genome sequencing data. The algorithm presented here can handle missing values and thus loci with incomplete data across samples do not need to be excluded.

### Preprocessing

`asmultipcf` uses the same pre-processing steps as the allele-specific single sample algorithm proposed by Nilsen et al. (2012). Most importantly BAFs are mirrored in order to obtain a single track in regions of allelic imbalance and extreme outliers, which are likely to be noise, are removed from logR and BAF data (see Nilsen et al. (2012) for details of the pre-processing). Given the input data of *n* samples across *p* SNP loci, the pre-processing yields a single matrix **Y** = (**y**_*ij*_) ∈ ℝ^2*n×p*^ that contains both logR and BAF values.

### An exact algorithm for weighted segmentation

We extend the penalized least squares approach of Nilsen et al. (2012) to evaluate the fit of a segmentation solution to the data, and use a weighted least squares function to model missing values in the data matrix. A weight matrix **W** = (**w**_*ij*_) ∈ ℝ^2*n×p*^ is derived by assigning *w_ij_* a weight of 0 if *y_ij_* is missing and 1 otherwise. Then all missing values in **Y** are assigned an arbitrary (non-NA) value. Our aim is to find a segmentation *S* = *{I*_1_, *…, I_M_}* that minimizes the cost function

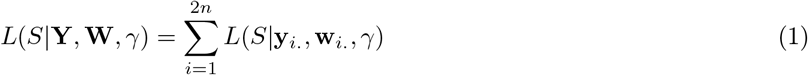

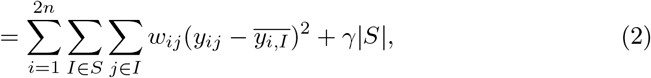

where the best fit on a given segment *I* is the weighted average of the observations on that segment

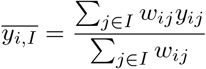

and where *γ* is a penalty parameter that controls the number of segments. Expanding the square in (2) and omitting the term that is independent of the segmentation we find

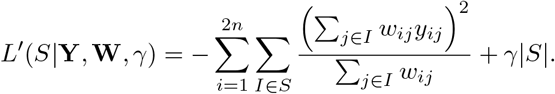

To find an optimal solution to the cost function we adapt the dynamic programming algorithm presented by Nilsen *et al*. (2012) to our weighted problem.

### Algorithm 1: asmultipcf

*Input:* Matrix **Y** of log-transformed copy numbers and B allele frequencies; Weight matrix **W**; penalty *γ >* 0;

*Output:* Segment start indices and segment averages

1. Calculate scores by setting **A**_0_ = [], **C**_0_ = [], **e**_0_ = 0 and iterate for *k* = 1, …, *p*

- **A**_*k*_ = [**A**_k-1_ 0] + **w**_.*k*_**y**_.*k*_
- **C**_*k*_ = [**C**_k-1_ 0] + **w**_.*k*_
- **d**_*k*_ = −1^*T*^(**A**_k_ ◦ **A**_k_ ◦ 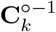) where ◦ denotes an element-wise matrix product and 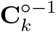 the element-wise inverse
- e_*k*_ = [e_*k*-1_ min(**d**_*k*_ + e_*k*-1_ + γ)]

storing also the index *t_k_* ∈ 1, …, *k* at which the minimum in the last step is achieved.
2. Find segment start indices from right to left as *s*_1_ = *t_p_*, *s*_2_ = *t_s_*_1 *−*1_, *…*, *s_M_* = 1, where *M* ≤ 1.
3. Find segment averages

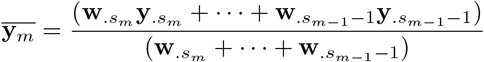

### A heuristic algorithm for large data sets

Algorithm 1 is of order *O*(*p*^2^), which means that the segmentation becomes computationally expensive for long sequences. However, instead of allowing breakpoints at any of the *p* positions, we can pre-select potential breakpoints and thereby reduce the runtime to *O*(*q*^2^) where *q* is the number of potential breakpoints. To identify potential breakpoints, different heuristics can be used. Here, we apply Algorithm 1 to overlapping subsequences, combine all of the inferred breakpoints and use them as input for the subsequent global segmentation. As in the implementation by Nilsen *et al*. (2012) we use subsequences of length 5000 with an overlap of 1000. Algorithm 2 describes the fast heuristic version of `asmultipcf`.

### Algorithm 2: Fast asmultipcf

*Input:* Matrix **Y** of log-transformed copy numbers and B allele frequencies; Weight matrix **W**; penalty *γ >* 0;

*Output:* Segment start indices and segment averages

1. Split data set into overlapping subsequences and apply steps 1 and 2 of Algorithm 1 to each of them in order to find potential breakpoints *r*_0_, *r*_1_, *…*, *r_q_* where *r*_0_ = 1 and *r*_1_ = *p* + 1.
2. Aggregate sequences between breakpoints by setting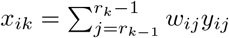and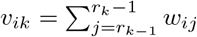.
3. Calculate segmentation solution by using the aggregated matrices **X** and **V** R^2*n×q*^ as input to Algorithm 1 instead of **Y** and **W**, respectively.

### Post-processing

Both algorithms yield a single segmentation solution *S* for all samples. However, we expect that only some of the segments will be shared between all samples while others will be private. While ASCAT can be run directly on the global segmentation solution, removing unnecessary breakpoints on a per sample base can reduce noise in the segment average estimates by generating larger segments. To refine breakpoints individually for each sample, we simply use the breakpoints inferred from the multi-sample segmentation and rerun steps 2 and 3 of Algorithm 2 on each sample individually based on these potential break points.

### Implementation

`asmultipcf` is part of the ASCAT R package from version 2.5 onwards. The asmultipcf function contains a parameter to select whether the exact or the fast algorithm should be run, as well as an option to include the per-sample breakpoint refinement. Furthermore, samples can be weight adjusted to account for quality differences in the data. An example of how to use this function can be found in the package manual.

## 3 Discussion

The independent segmentation of related samples can artificially inflate tumor heterogeneity. The algorithm presented here addresses this problem by joint segmentation. While this approach can potentially underestimate tumor heterogeneity, because CNAs that are shared by many samples are more likely to be detected than CNAs that are private or shared by only few samples, in practice the penalty parameter *γ* can be adjusted to ensure sensitivity. Overall, `asmultipcf` substantially improves the analysis of copy number changes of multiple samples.

## References

Mangiola, S. et al. (2016). Comparing nodal versus bony metastatic spread using tumour phylogenies. Scientific Reports, 6, 33918 EP –.

Nilsen, G. et al. (2012). Copynumber: Efficient algorithms for single-and multi-track copy number segmentation. BMC Genomics, 13(1), 591.

Schwarz, R. F. et al. (2015). Spatial and temporal heterogeneity in high-grade serous ovarian cancer: A phylogenetic analysis. PLOS Medicine, 12(2), 1–20.

Van Loo, P. et al. (2010). Allele-specific copy number analysis of tumors Proceedings of the National Academy of Sciences, 107(39), 16910–16915.

